# Robustness Revisited: On the Neutral Evolution of Centrality and Localization

**DOI:** 10.1101/2024.08.29.610396

**Authors:** Yehonatan Sella, Aviv Bergman

## Abstract

This study investigates the intricate interplay among neutral landscape structure, mutation rate, recombination rate, and population dynamics in shaping evolutionary robustness. We provide a comprehensive framework that elucidates how different evolutionary forces interact to influence genotypic robustness and localization within haploid and diploid populations. We demonstrate that in haploid populations, high mutation rates relative to recombination typically drive the population toward regions of increased eigencentrality, a graph-theoretic measure of centrality which is correlated while not identical to mutational robustness. On the other hand, recombination increases the localization of the population to a smaller region of genotypic space, while high values of recombination relative to mutation can introduce shifts in distribution away from eigencentrality and toward attractors of the recombination dynamics. Diploid dynamics further complicate these interactions, showing reduced alignment with eigencentrality under both high mutation and recombination rates, with the exception of structured diploid landscapes where dynamics are still aligned with increasing eigencentrality. Our findings underscore the nuanced dependencies of evolutionary outcomes on both local and global landscape structures as well as evolutionary parameters.

**Significance Statement:** Our work advances the theory of neutral evolution, paying particular attention to the question of how the holistic fitness landscape structure shapes the process of evolution and gives rise to emergent evolutionary phenomena. Since neutral evolution does not depend on direct selection, its ramifications can be both subtle, as they depend on network-wide properties, and ubiquitous, as they are not tied to context-specific adaptations. Our study provides a theoretical framework that connects the structure of neutral fitness landscapes with the dynamics of mutation and recombination rates, and the distinct behaviors of haploid and diploid populations. We establish general heuristic principles regarding the way evolutionary outcomes, such as robustness and localization, are influenced by the interplay of these factors.

## Introduction

The evolution of genetic robustness has long fascinated evolutionary biologists. Robustness, originally termed by Waddington “canalization”, was introduced to account for “the very general observation … that the wild type of an organism, that is to say, the form which occurs in Nature under the influence of natural selection, is much less variable in appearance than the majority of the mutant races” [1]. Ever since then, numerous studies took upon themselves to corroborate and expand on this phenomenon [2, 3]. While robustness has often been studied under the lens of adaptive evolution and selective pressures, a growing body of work suggests that neutral evolutionary processes—where most mutations are either neutral or slightly deleterious—can also play a significant role in shaping robustness. Specifically, more recent studies, incorporating development in the context of gene regulatory networks, have shown that robustness can evolve at the absence of stabilizing selection and that selection for developmental stability suffices to induce phenotypic robustness [4, 5, 6, 7].

The importance of studying evolution in the absence of stabilizing selection is underscored by Kimura’s neutral theory of evolution [8], which posits that, at least at the biomolecular level, most mutations are either neutral or deleterious, and that it is these neutral mutations which are responsible for most gene substitutions in a population.

The framework of neutral evolution can be profitably combined with the powerful metaphor of fitness landscapes introduced by Sewall Wright in the early 1930’s and later revisited in the late 1980’s [9, 10, 11]. A fitness landscape refers to a network of potential genotypes for a population, connected via mutation, together with a fitness value assigned to each genotype. The metaphor is suggestive of a population that walks along the landscape of genotypes in a direction of increasing fitness, climbing toward a peak [12]. However, in the neutral setting, the idea of climbing toward peaks is de-emphasized given the flat nature of the landscape. Neutral landscapes, characterized by a set of genotypes that all share a similar fitness level, provide a unique perspective on evolutionary dynamics. Gavrilets [13, 14, 15], who studies such landscapes at length, has evocatively termed them “holey landscapes” to emphasize that they can be viewed as flat surfaces riddled with holes of lower fitness, and that the imperative of evolution is not so much to climb to higher peaks but to avoid falling in the holes. In these landscapes, the movement of populations is not solely directed by fitness gradients but rather influenced by the connectivity of genotypic networks and the interplay of evolutionary mechanisms like mutation and recombination. Understanding how these factors interact is crucial for comprehending the evolutionary pathways that lead to robustness.

Previous studies have begun to chart the implications of evolution on neutral landscapes. A key paper [16] has shown that evolution of large populations on neutral landscapes can be well-modeled by assuming for simplicity that all deleterious mutations are fatal, so that the fitness is either 0 or 1, allowing us to restrict our attention to the set of neutral genotypes with fitness 1. Under this model, assuming infinite haploid populations and only mutation (no recombination), the population will always converge to a distribution described by the unique leading eigenvector of the adjacency matrix of the graph of viable genotypes, as long as this graph is connected. This convergence is independent of the choice of mutation rate or of the initial population distribution. Given the importance of this fact, we include a separate proof in the supplemental information. Moreover, the *eigenvalue* of this matrix measures the average mutational robustness of the stationary distribution.

The unique leading eigenvector of a graph’s adjacency matrix is known as its *eigencentrality*. Its uniqueness (up to scaling) is guaranteed by Perron-Frobenius [17], which also ensures it is the unique eigenvector all of whose entries are positive.

Eigencentrality is a well-known graph-theoretic measure of the “centrality” or “influence” of a node in a graph. That is, for a given vertex *v*, we can interpret the *v*th component of the eigencentrality vector as a score, where a higher score indicates greater eigencentrality. Letting *λ* ∈ ℝ be the top eigenvalue of *A*, and letting **ec** denote the leading eigenvector, the eigenvector equation *A***ec** = *λ***ec** means that for each vertex *v* in the graph,

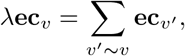

where *v’* ∼ *v* represents the set {*v’*} of one-step neighbors of *v*. Thus eigencentrality rewards a node for having many neighbors, but only if those neighbors themselves are eigencentral.

The observation that the frequency of a genotype in the stationary distribution is precisely given by that genotype’s eigencentrality thus formalizes the intuition that evolution drives the population to central regions of the landscape, away from the holes.

Centrality is closely related to robustness. In a neutral landscape, one may define the mutational robustness of a genotype as the fraction of its mutational neighbors that are viable, which is proportional to its degree in the graph of viable genotypes. While degree and eigencentrality can markedly diverge depending on the structure of the graph, eigencentral nodes tend to have large degree, thus explaining the evolution of robustness on neutral landscapes under mutational dynamics. We further explore the interplay between different measures of centrality on different families of landscapes in the present paper.

A notable property of eigencentrality on graphs that are sufficiently irregular, to be discussed further in the main body of the paper, is *localization*. Namely, for these networks, high eigencentrality scores are limited to a small set of nodes which are close together [18, 19, 20], while exponentially decaying away from this localized region. The precise degree of localization depends on the structure of the network. This property is noteworthy since it suggests that, even under the paradigm of neutral evolution, the course of evolution is not entirely diffuse, but may be steered to a particular localized region of genotypic space.

While the stationary distribution under mutation dynamics has a simple analytical description in terms of eigencentrality, once recombination is introduced the dynamics become more complicated, and the stationary distribution depends on the balance between the rates of mutation and recombination, as well as on the specific initial distribution. A recent paper [21] analyzed the mutation-recombination dynamics on holey landscapes, concluding that recombination tends to further increase population robustness. Their paper also suggests the importance of a recombination weight in determining the most frequent genotype in the stationary distribution, when recombination dominates over mutation.

In this paper, we further elucidate and complicate these observations. Specifically, we show that while mutation tends to drive populations toward regions of high eigencentrality, indicating robustness to genetic perturbations, recombination can either enhance or disrupt this robustness depending on the underlying landscape structure and population dynamics. We place particular emphasis on studying localization along with robustness and its more refined counterpart, eigencentrality. We show that increased recombination consistently leads to increased localization of the stationary distribution, but that the impact of recombination on robustness is more complicated. We show that, for certain families of landscapes, there are distinct phase shifts associated with the balance between mutation and recombination: if mutation is large enough compared to recombination, then the stationary distribution resembles that of eigencentrality, only more localized, whereas if mutation falls below a certain threshold, the stationary distribution often shifts to a different part of the graph, which partly correlates with eigencentrality and robustness, but much less reliably so. We separate the nature of increased population robustness into two distinct phenomena: increased localization, and increased robustness (or eigencentrality) of the mode. While recombination reliably increases the former, mutation more reliably increases the latter.

We also systematically study the influence of the initial population on the stationary distribution. In addition, we study the dynamics under both haploid and diploid models, and show that these dynamics substantially differ under the two models, with eigencentrality lower in diploid populations even when mutation is high relative to recombination, unless the fitness landscape is specially structured, as discussed further below.

### Evolution on abstract fitness landscapes

We describe the process of evolution on fitness landscapes abstractly, in terms of both mutation and recombination. We define a fitness landscape to be the data of a graph *G* = (*V, E*) together with a fitness function *w* : *V* → ℝ^≥0^ valued in non-negative numbers. For simplicity we assume the graph is undirected and unweighted. We will further make the simplifying assumption that the graph *G* is regular, so that each vertex has the same degree *N* for some *N* ≥ 1.

Each vertex *v* ∈ *V* stands for an entire genotype, a point in genotypic space. There is an edge *v*_1_ ∼ *v*_2_ between two vertices *v*_1_, *v*_2_ ∈ *V* if a single mutation would transform *v*_1_ to *v*_2_ and vice versa. The fitness function *w*(*v*) describes the probability of an individual with genotype *v* surviving to reproduce. We assume uniform fecundity for all genotypes.

The standard biological example to keep in mind is the graph of all sequences of length *n* over a given alphabet *A*, where two sequences are neighbors if there is a single edit that transforms one to the other. In the case that *A* is an alphabet of size two, this graph is the *n*-dimensional hypercube, denoted *C*_*n*_. However, for the sake of abstractness and generality, we do not restrict ourselves to this biological case.

Define a node *v* to be *viable* if *w*(*v*) *>* 0 and *inviable* if *w*(*v*) = 0. Since inviable nodes are thrown out by evolution, it is useful to consider the restricted graph *G’* = (*V’, E’*), where *V ’* ⊂ *V* consists only of the viable nodes, and *E*^*i*^ is the restriction of *E* to *V* ^*i*^. We will make the assumption that the graph *G’* is connected. If it had multiple connected components, one can study them separately since they never interact, at least assuming no recombination and only one-step mutations.

Under this simple abstract framework, we can model evolution as a stochastic (Markovian) process on this graph *G’*. We begin with the case of asexual reproduction, and later treat the more involved case of sexual reproduction. We also make the simplifying assumption of a very large population, so that we can ignore stochastic effects.

Let **x**_*t*_ denote the expected population distribution at time *t*, which we think of as a vector in ℝ^|*V ’*|^; that is *x*_*t,v*_ is the expected number of individuals with genotype *v* at time *t*. Now, let **p**_*t*_ denote the probability distribution of the population at time *t*, which is simply a normalized version of **x**_*t*_, namely, we have 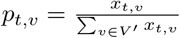.

### Mutational dynamics

We first model evolution with the following asexual model, which depends on the specification of a mutation rate 0 *< µ <* 1. Each individual with genome *v* has probability *w*(*v*) of surviving and reproducing. If it survives, there is a probability (1 − *µ*) that the offspring’s genotype is still *v*, and a probability of *µ* that it has incurred a mutation. In this event, the new genotype is randomly, uniformly selected from among the neighbors of *v* in the graph *G*. The mutant is deleted if it is inviable.

The evolutionary process is specified by recurrence equations for **x**_*t*+1_ in terms of **x**_*t*_, as follows:

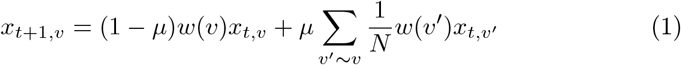

We can rewrite this as a matrix equation. Let *A* be the |*V’*| *×* |*V’*| matrix recording the edges *E’* between *viable* nodes. That is, we let *A*_*vw*_ = 1 if *v, w* are neighbors which are both viable, and *A*_*vw*_ = 0 otherwise. Let *W* be the |*V ’*| *×* |*V ’*| diagonal matrix with *W*_*vv*_ = *w*(*v*). Then we have

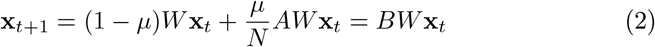

for 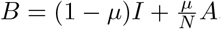.

Note that the vectors **x**_*t*+1_ will decay exponentially since each individual produces on average less than one offspring. However, we are implicitly assuming a population whose size approaches infinity, and are therefore more interested in the evolution of the normalized vectors **p**_*t*_ describing the probability distribution of the population among the viable nodes.

### Reproductive dynamics

Next, we describe sexual reproduction dynamics abstractly. To this end, the graph *G* = (*V, E*) must be augmented with the additional structure of a reproduction tensor, which is a function *R* : *V × V × V* → ℝ^≥0^ symmetric with respect to *v*_1_ and *v*_2_.

The number *R*(*v*_1_, *v*_2_, *v*_3_) is interpreted as the probability that an offspring with genotype *v*_3_ is produced from a mating of parents with genotypes *v*_1_ and *v*_2_ (discounting mutation). In practice, we only require the information of this tensor on viable nodes; let *R’* : *V’ × V’ × V’* → ℝ^≥0^ be the restriction of *R* to *V ’*.

The tensor *R* will depend on the rate of recombination. To elucidate this, let 0 ≤ *r* ≤ 1 be the recombination rate. Reproduction can be conditioned on the presence or absence of a recombination (crossing-over) event. To represent these two cases, let *R*_0_ : *V × V × V* → ℝ^≥0^ be a tensor which describes reproduction without recombination, and *R*_1_ : *V × V × V* → ℝ^≥0^ be a tensor which describes reproduction with recombination (here we restric the process to a single recombination event), so that *R*_0_(*v*_1_, *v*_2_, *v*_3_) (respectively *R*_1_(*v*_1_, *v*_2_, *v*_3_)) represents the probability that if parents with genotypes *v*_1_ and *v*_2_ mate without (respectively, with) recombination, they produce an offspring with genotype *v*_3_.

Then the total reproduction tensor may be written as

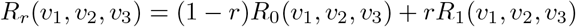

The subscript indicates the dependence on the parameter *r*. The tensors *R*_0_ and *R*_1_ are assumed to both be stochastic in the third variable, in the sense that if *v*_1_, *v*_2_ are fixed, and for a given *i* = 0, 1, the probabilities *R*_*i*_(*v*_1_, *v*_2_, *v*_3_) sum to 1. If the tensor is restricted to *V* ^*i*^ it will be substochastic in the third variable.

Finally, we define a symmetric quadratic function *Q* : ℝ^|*V*^*’*^|^ → ℝ^|*V*^*’*^|^ by

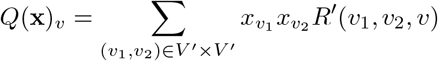

Reproduction dynamics can then by defined simply by **x ⟼** *Q*(**x**). If we interpret **x** as a population count vector, the dynamics **x ⟼** *Q*(**x**) reflect a panmictic population in which every individual mates with every other individual in the population to produce the next generation. That is, we are assuming infinite population and are primarily interested in tracking the population distribution rather than absolute numbers; the resulting dynamics on population distribution captures random mating.

Combined with mutation and selection, the process becomes

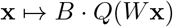

The standard biological example is that of either haploid or diploid populations, and in which the recombination scheme is given by a set of recombination events. Below we will study both cases of haploid and diploid populations.

In describing a function *R* : *V × V × V* → ℝ, it will be convenient to recast it as a function *R* : *V × V* → ℝ^|*V* |^, where 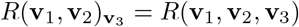.

In the haploid case, *G* = (*V, E*) is the *n*-dimensional hypercube graph *C*_*n*_.

Here 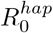 represents randomly choosing one of the parents, 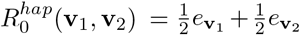, where *e*_**v**_ denotes the standard basis vectors. We focus on a recombination scheme that assumes one crossover event, where the crossover locus is picked at random from among the *n* − 1 possibilities. This is described by the recombination tensor 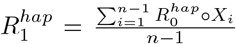, where *X*_*i*_ : *C*_*n*_ *× C*_*n*_ → *C*_*n*_ *× C*_*n*_ is the crossover operation at the *i*th locus, i.e. *X*_*i*_((*a*_1_, …, *a*_*n*_), (*b*_1_, …, *b*_*n*_)) = ((*a*_1_, …, *a*_*i*_, *b*_*i*+1_, …, *b*_*n*_), (*b*_1_, …, *b*_*i*_, *a*_*i*+1_, …, *a*_*n*_)). As above, we then have 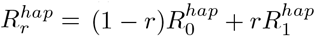.

In the diploid case, rather than proceeding by specifying *R*_0_ and *R*_1_, we relate it to the haploid case using the following general construction: given any set *V* and tensor *R* : *V × V × V* → ℝ, it induces a tensor on *V × V*, that is, a map *R × R* : (*V × V*) *×* (*V × V*) *×* (*V × V*) → ℝ by multiplication, that is ((**v**_1_, **v**’_1_), (**v**_2_, **v**’_2_), (**v**_3_, **v**’_3_)) ⟼ *R*(**v**_1_, **v**_2_, **v**_3_)*R*(**v**’_1_, **v**’_2_, **v**’_3_).

We then define 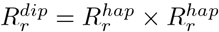 as a tensor on *C*_*n*_ *× C*_*n*_.

Note that different values of *r* will be appropriate depending on the biological context. For example, if the graph *V* represents nucleotide sequences in a small contiguous subset of the total genome, then a small value of *r* is reasonable, given that a crossover event is unlikely to happen within this small subset of the genome. On the other hand, if *V* represents the total genome, or if the loci represent genes rather than nucleotide base pairs, then a larger value of *r* is appropriate. (Consider moving to discussion)

### Holey landscapes

We will primarily focus on the case of “holey” landscapes. Namely, those in which all viable individuals have the same fitness (for simplicity we set it at 1, though that does not affect our results), and so the only fitness distinction is between viable and nonviable genotypes.

In the case of holey landscapes, the matrix *W*, which records the fitness of viable genotypes, is the identity. Thus the mutation process becomes

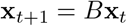

And together with recombination, it becomes

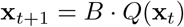

Crutchfield et al’s result [16], mentioned in the introduction, that mutational dynamics converge to eigencentrality, follows from the fact that the eigenvectors of *B* are the same as those of *A*, together with the Perron-Frobenius theorem. See the Supplemental Information for a proof.

### Measures of centrality, robustness and localization

Next we study how robustness relates to eigencentrality. On holey landscapes, we equate viability with phenotype. The robustness of a genotype is then the proportion of its mutational neighbors which are viable. As a vector equation, the robustness vector **r** ∈ ℝ^|*V*^’^|^, where **r** is the robustness of *v* ∈ *V ’*, is given by 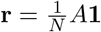. Here **1** is the vector of ones.

We can generalize this notion of robustness to define the *k*th-order robustness of a node *v* ∈ *V ’* as 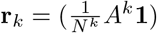. The number *r*_*k*_(*v*) measures the probability, given a path of length *k* in the graph *G* starting at *v* (representing a sequence of mutations), that all the nodes along the path are viable. The vector **r**_*k*_ records these probabilities across all *v*. Of course, **r**_1_ equals the above measure of robustness **r**, and the **r**_*k*_, after renormalizing, converge to eigencentrality **ec**. We can integrate the different *k*th-order robustnesses in one measure by computing the average length of a random walk in *G* before reaching an inviable node. That is, for each *v* ∈ *V ’*, consider the random walk which starts at *v* and where each time, the next node is chosen uniformly at random from among the neighbors of *v* in *V*. The random walk is terminated as soon as an inviable node is visited. let *L*_*v*_ be the random variable measuring the length of this random walk, that is, the length of the path taken in *V ’*, not including the terminal inviable node. Then define *l*(*v*) = *E*(*L*_*v*_), the expected length of this random walk. A low value of *l*(*v*) would indicate that *v* is precariously located on the edge of the region of viability, while a high value of *l*(*v*) would indicate *v* is securely within the region of viability. Let **l** denote the vector recording *l*(*v*) across all *v* ∈ *V ’*.

The measure *l*(*v*) is related to the *k*th-order robustnesses by the formula 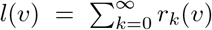, where we define *r*_0_(*v*) = 1. Indeed, this follows from the fact that *r*_*k*_(*v*) = *P* (*L*_*v*_ ≥ *k*). The vector **l** can then be calculated by 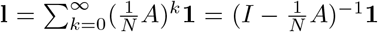. Note that the matrix 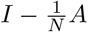 is invertible as long as *N* is not an eigenvalue of *A*, which it is not unless all states are viable.

We can obtain expressions for the average values of **r**_**k**_ and **l** across the stationary population distribution, under mutational dynamics. Let 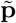 be the stationary distribution of evolution, which we know is the eigencentrality vector, normalized to sum to 1. Crutchfield et. al have shown that the average population robustness, that is, 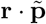, is equal to 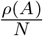, where *ρ*(*A*) is the specral norm of *A*, which is the top eigenvalue of *A*. More generally, it is a simple derivation that the average population *k*th-order robustness, given by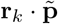, equals 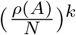.

Indeed, substituting in the equation 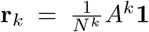, we see that 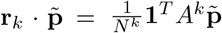 (since *A* is symmetric). But 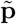 is an eigenvector of *A* with eigenvalue *ρ*(*A*). So 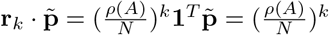.

The average value of **l** across the stationary distribution **p** is then 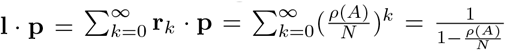. Note that *ρ*(*A*) ≤ *N* always, the case *ρ*(*A*) = *N* occuring only if all states are viable.

Next we argue that evolution under mutation tends to favor robust nodes, whether this robustness is measured by *r*(*v*), *r*_*k*_(*v*) or *l*(*v*). We can express this fact in two different, but equivalent ways:

First, that the average population robustness of the stationary distribution will always be greater than the population robustness of the uniform distribution, representing a “random” distribution; Second, that the correlation between the robustness of a genotype and its frequency in the stationary distribution is positive.

Indeed, the uniform distribution is given by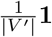, and so the average *k*thorder robustness of the uniform distribution is 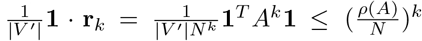, where we use the fact that the number **1**·*A*^*k*^**1** is bounded by *ρ*(*A*)^*k*^||1||^2^ = *ρ*(*A*)^*k*^||*V’* || (see Supplemental Information for more details). Thus, the average *k*th-order robustness of the uniform distribution is bounded above by that of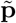.

Equality is only obtained if the uniform distribution 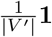 already equals the stationary distribution 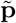. In this case, all viable nodes have the same number of viable neighbors, and so robustness is uniform across all nodes.

One can recast this calculation as showing that the correlation between the stationary distribution and robustness is positive. Indeed, the difference 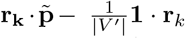, which we have shown is positive, is precisely 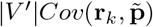.

As a consequence, we also see that the stationary distribution 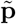 positively correlates with **l**, since the latter is a positive combination of the **r**_*k*_.

While we have shown in general that 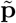 correlates positively with the measures **r, r**_*k*_, **l**, the precise extent of these correlations varies and depends on the specific structure of the network of viable nodes.

### Localization

A notable property of eigencentrality in certain landscapes (or graphs) is localization. Namely, high eigencentrality scores are limited to a small set of nodes which are close together [18, 19, 20].

The precise degree of localization depends on the structure of the network. To measure localization in a given population distribution **p**, One can consider the use of two measures:

First, the Simpson diversity index *SD*(**p**) = Σ_*v*∈*V ’*_ (**p**_*v*_)^2^ = **p** · **p**, which measures the probability that two randomly chosen individuals in the population share the same genotype. We have 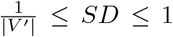, where a larger value of *SD* indicates greater localization, the minimum being attained at a uniform distribution, and the maximum of *SD* = 1 attained at an isogenic distribution. Secondly, one can measure the expected distance (shortest-path distance in the graph *G*, which translates to Hamming distance in the biological setting) between two randomly chosen individuals, 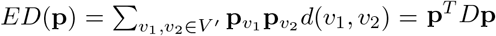, where *D* is the |*V ’* |*×*|*V ’* | distance matrix. A lower value of *ED* indicates increased localization, with *ED* = 0 for isogenic distributions, and where *ED* is bounded above by the diameter of the graph.

Normalized versions of these localization measures, in which we normalize by the localization of the uniform distribution, can also be used: 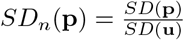 and 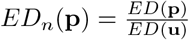, where **u** denotes the uniform distribution on *V ’*.

Below we investigate the above measures of robustness and localization for stationary distributions on specific families of holey landscapes (to illustrate our results, for the localization measure, we employed the Simpson diversity measure).

### Families of holey landscapes

We present a few examples of families of holey landscapes, which we will study throughout this paper.

#### Example 1: Geometric examples

To provide geometric intuition, we first provide examples of holey landscapes on geometric grids. Though these are not directly applicable to networks of genomes, they illustrate the concepts under discussion and may be easily visualizable for 1and 2-dimensional grids. Moreover, these graphs can in principle be embedded as subgraphs of hypercubes and thus attain a genetic context.

##### Example 1.1: Box in ℤ^d^

Consider the *d*-dimensional grid ℤ^*d*^, with the edge relation given by adjacency on the grid. Let *n*_1_, …, *n*_*d*_ be positive numbers, and say that a vertex (*a*_1_, …, *a*_*d*_) is viable if 1 ≤ *a*_*i*_ ≤ *n*_*i*_ for all *i*.

##### Example 1.2: Russian Roulette on a grid

The Russian Roulette holey landscape, so-named by Gavrilets [13, 14, 15, 22], is a grid as in Example 1.1, in which nodes have a certain independent probability *p* of being viable. This process of removing points at random from a network is also nown as *site percolation* in percolation theory.

Percolation theory studies the connected components of the resulting graph in the infinite setting of ℤ^*d*^ as opposed to finite grids. In particular, there is a critical threshold *p*_*c*_ such that if *p > p*_*c*_, the resulting graph has an infinite connected component, whereas if *p < p*_*c*_, then the resulting graph has no infinite connected components (almost surely). For example, for Z^2^, the critical probability is known to be approximately *p*_*c*_ ≈ 0.593 [23]. We will employ values of *p* larger than *p*_*c*_ in order to obtain large connected components, and we will focus on the largest connected component.

### Example 2 : Genetic examples (hypercube subgraphs)

While geometric examples, in particular in two or three dimensions, can help by providing visualizable examples, this geometric intuition can be misleading, as Wright himself observed about his concept of the fitness landscape [9, 10]. It is therefore best to check any intuition gained by the geometric examples against landscapes that are grounded in the genetic context, namely subgraphs of the hypercube {0, 1}^*n*^.

#### Example 2.1 : Mesa landscapes

A Mesa landscape is specified by a dimension *n*, a choice of central node *v* ∈ {0, 1}^*n*^ and a threshold number *a* where 0 *< a < n*. A node *v*^*i*^ ∈ {0, 1}^*n*^ is then viable if its Hamming distance from *v* is at most *a*. Biologically, viability is maintained as long as the genotype lies within a certain threshold of an “optimal” genotype. By symmetry, we can always assume for simplicity that *v* is the string (0, 0, …, 0), so that a node is viable if the number of ones in the corresponding binary is bounded above by *a*.

Mesa landscapes are in some sense analogous to the boxes of Example 1.1, in that there is a clear central node, and a clear boundary lying a given distance from that central node.

#### Example 2.2: Genetic Russian Roulette

In the genetic Russian Roulette, each node in the hypercube {0, 1}^*n*^ has independent probability *p* of being viable. We then take the largest connected component.

### Eigencentrality, robustness and localization for mutation-only dynamics

We now investigate the properties of the stationary distribution of mutational dynamics– that is, eigencentrality– on the above families of landscapes. In this section we will denote the eigencentrality distribution by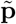.

First, for the sake of visualization, we plot eigencentrality for representative landscapes of each of the families described above, see Figure 1.

**Figure 1:**
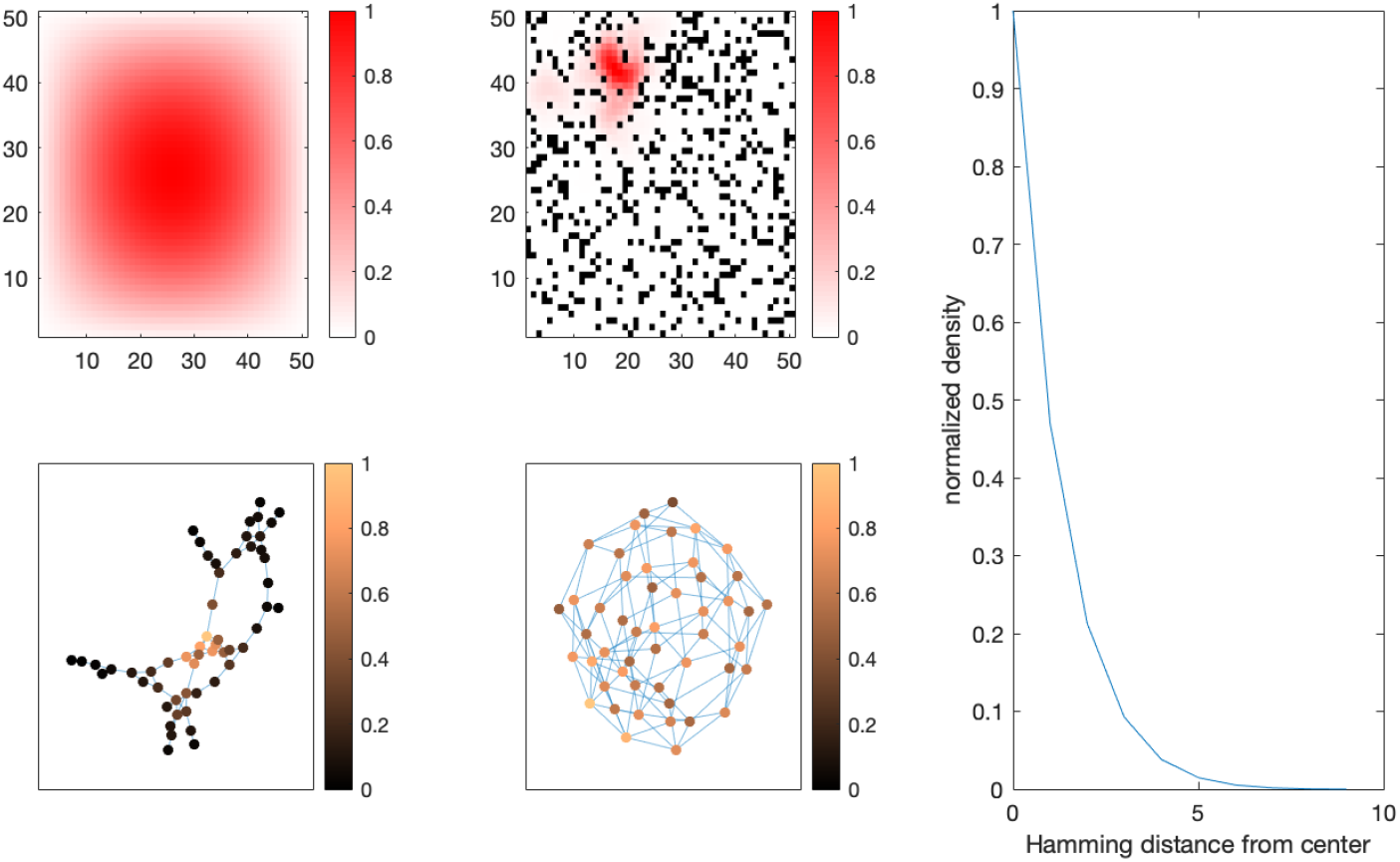
Eigenplot. Plotting the eigencentrality distribution for representative landscapes of different types: a) a 2-dimensional geometric grid with boundary interpreted as inviable, and mutations being neighbors on the grid. b) a 2-dimension geometric landscape with holes. c) a genetic landscape with low probability of viability. d) a genetic landscape with high probability of viability. e) a “Mesa” landscape (genetic landscapes where a genotype is viable as long as it is within a certain Hamming distance of the “optimal” genotype. Note that landscapes a) and b) serves only for geometric intuition, not being biologically motivated, while c) and e) are more biologically relevant. The phenomenon of localization is illustrated in examples b) and c), where we see that the high-eigencentrality portion of genotypic space is limited to a small area, rather than being diffuse. The other examples are not characteristic of localization.

In the geometric box as well as the Mesa landscape, there is a clearly defined center, and eigencentrality peaks at that center and diminishes away from the center, approaching 0 toward the boundary. In the geometric case of an *n*_1_ *×* · · · *× n*_*d*_ box, one can determine analytically that the stationary distribution is (up to rescaling)

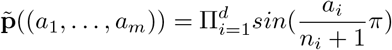

(See Supplemental Information for a proof).

On the other hand, for the geometric and genetic Russian Roulette families, there is no clear center to the landscape. In the geometric Russian Roulette landscapes in two dimensions in which *p* is not too close to 1, eigencentrality is localized to a small region of the space, a pocket which is a relatively high-viability region. Similar localization is observed for the genetic Russian Roulette if *p* is small. However, for larger *p* eigencentrality is more diffuse.

To support these results systematically, we calculate the localization measures *SD*_*n*_ for these families of landscapes over varying parameters. The following image shows normalized Simpson diversity (*SD*_*n*_) for different families of landscapes, under varying parameters.

For the stochastic families, we generated 50 landscapes for each parameter choice, and took the mean *SD*_*n*_ across these landscapes. In the geometric Russian roulette, we see that a lower probability of viability increases localization, as does increased grid size. Similarly in the genetic Russian roulette, localization tends to increase with lower probability of viability and higher dimension. The Mesa landscape serves as a control. We do not consider the Mesa landscapes as exhibiting localization, since the population is diffused throught the entire landscape as there are no internal barriers to diffusion. Indeed, while *SD*_*n*_ for the Mesa landscapes increases with dimension, it is still only around twice the base-level *SD* of a uniform distribution, whereas the Russian Roulette landscapes attain much higher values of *SD*_*n*_ for certain parameters.

As a control for the geometric landscapes, one can prove using an integral approximation (see Supplemental Information) that *SD*_*n*_ of an interval approaches 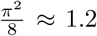 as the size of the interval approaches infinity, and *SD*_*n*_ of a square grid approaches 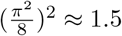 as the size of the grid approaches infinity.

Localization of eigenvectors of both the adjacency matrix and Laplacian of a graph is a well-studied phenomenon [19] have shown that graphs with more homogenous degree distributions tend to not exhibit localization, while graphs with more heterogenous degree distributions tend to exhibit localization, and moreover, the top eigenvector tends to be concentrated among a cluster of nodes with similarly high degrees. Indeed, the phenomenon of localization can be understood to emerge when a cluster of nodes with high degrees is surrounded by nodes with significantly lower degrees, which act as a barrier to diffusion.

These observations are consistent with our observations on the Russian roulette landscapes, since a lower probability *p* of viability entails a higher variance in the degree distribution relative to the mean degree (see Figure 2).

**Figure 2:**
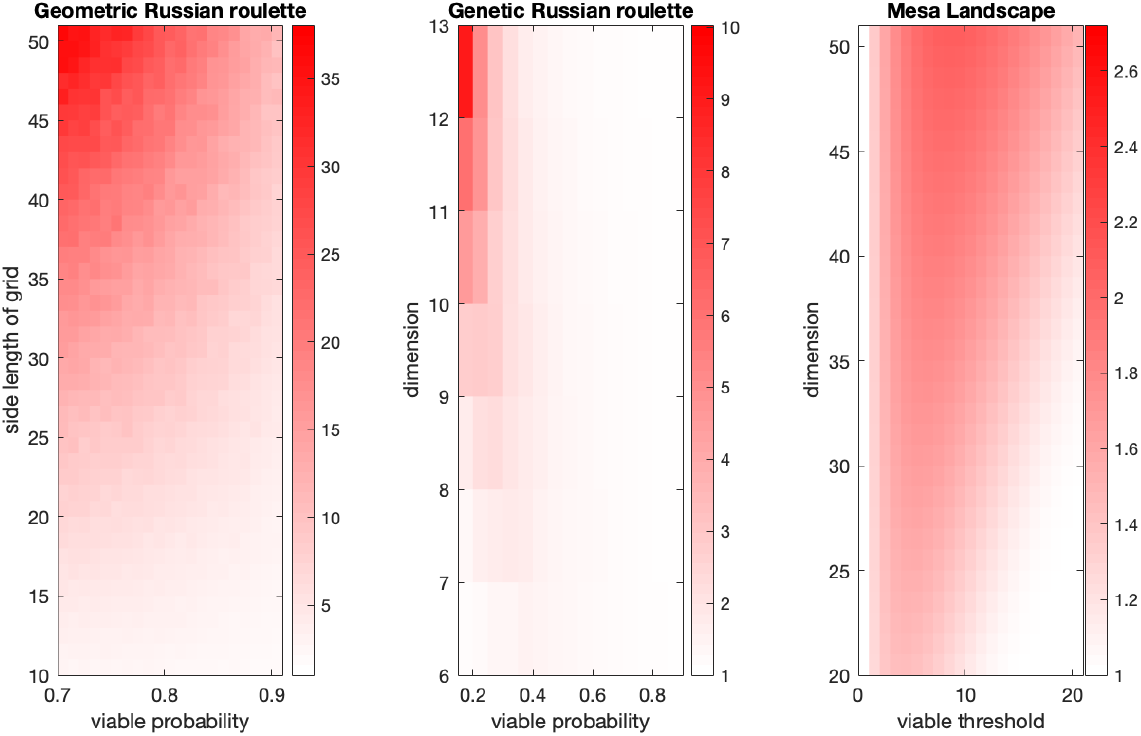
A systematic “heatmap” of the localization measure for different families of landscapes. This show the trend that a lower proportion of viable genotypes, as well as higher dimension, tends to increase localization, and that there seems to be a sharp transition from low-localization to high-localization regimes.

Next we consider robustness. As we have noted, average population robustness always increases upon evolution if the initial population distribution is uniform. We see below (Figure 3) that the increase is more pronounced, in both geometric and genetic versions of the Russian roulette landscapes, when the probability of viability is low, that is, when localization is present. This agrees with the intuition that a localized distribution is concentrated on a pocket of high-degree (i.e. high-robustness) genotypes, thus buffering against the low average robustness of the landscape as a whole. Non-localized eigencentrality distributions, while correlated with robustness, are diffused throughout the entire landscape. For another perspective on this observation, we plot eigencentrality (normalized such that the maximum is 1) of genotypes in a landscape against the number of viable neighbors, aggregated over 50 landscapes for each landscape type and visualized as box plots. We note that in the case of low probability of viability, exhibiting localization, the correlation between degree and eigencentrality is less predictable, with many outliers in which lower-degree genotypes have high eigencentrality and vice versa. In the case of high probability of viability, the correlation, though noisy, is much more straightforward.

**Figure 3:**
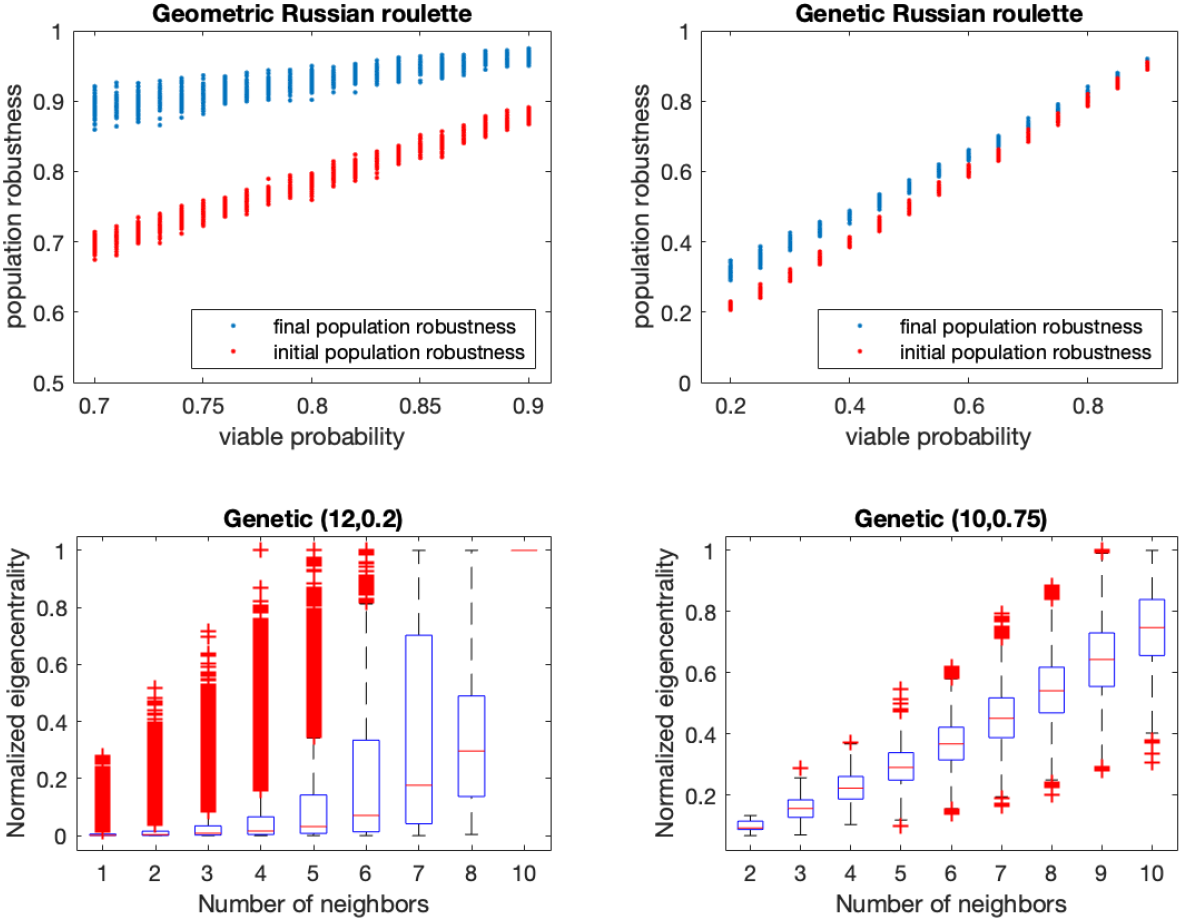
Robustness and Localization. The top two plots show the final versus initial measures of population robustness for random landscapes of both the geometric and genetic Russian roulette families of landscapes. We observe that in both cases, the trend is of increasing robustness over the course of evolution, while the magnitude of change is more pronounced in the geometric landscapes as well as when the viable probability is lower. The bottom plots show the distribution of normalized eigencentrality (normalized so that 1 is the maximum), broken down by a node’s number of viable neighbors. We see that the distributions have starkly different shapes between the low-*p* and high-*p* cases of genetic Russian roulette landscapes. In both there is a correlation between number of neighbors and eigencentrality, but in the low-*p* case there are many more outliers, nodes where the number of viable neighbors is relatively low while eigencentrality remains high. This is indicative of localization: being located in the right region of the network is more important than the immediate number of neighbors

### Eigencentrality, robustness and localization under mutation and recombination

We now explore the general differences between the dynamics of evolution under mutation versus recombination. We begin by studying the haploid case.

#### Haploid Recombination Dynamics

While asexual evolution on neutral landscapes reaches the same stationary distribution regardless of the mutation rate and the initial distribution, the stationary distribution reached by sexual reproduction with recombination depends on the rates of mutation and recombination as well as on the initial population distribution. In this section we investigate these dependencies and the relationship of the resulting stationary distribution to the eigencentrality distribution.

For the purposes of this section, we focus on genetic Russian Roulette landscapes, in both the low-*p* regime, where we see localization of eigencentrality, and in the high-*p* regime, where we do not. We collect data from 50 different randomly generated genetic Russian Roulette landscapes in each regime, and for each of these landscapes sweep the mutation rates between 0 and 0.5 (we sample more closely the smaller mutation rates, but include large ones for a full picture), while keeping recombination rate at 0.1. For the initial distribution, we focus on the two extremes: either a uniform distribution over the entire viable genotypic space, or an isogenic distribution at a single genotype.

The first observation is that the introduction of recombination increases the stationary distribution’s localization; the lower the rate of mutation is relative to recombination, the more pronounced this localization becomes. This qualitative relationship is observed regardless of *p* and regardless of the initial distribution, though it is more pronounced for low *p* and isogenic initial distribution.

As seen in Figure 4, the transition to high localization occurs rapidly as the mutation rate decreases past a certain inflection point. Thus, when the effect of recombination dominates over that of mutation, the resulting stationary distribution is substantially localized.

**Figure 4:**
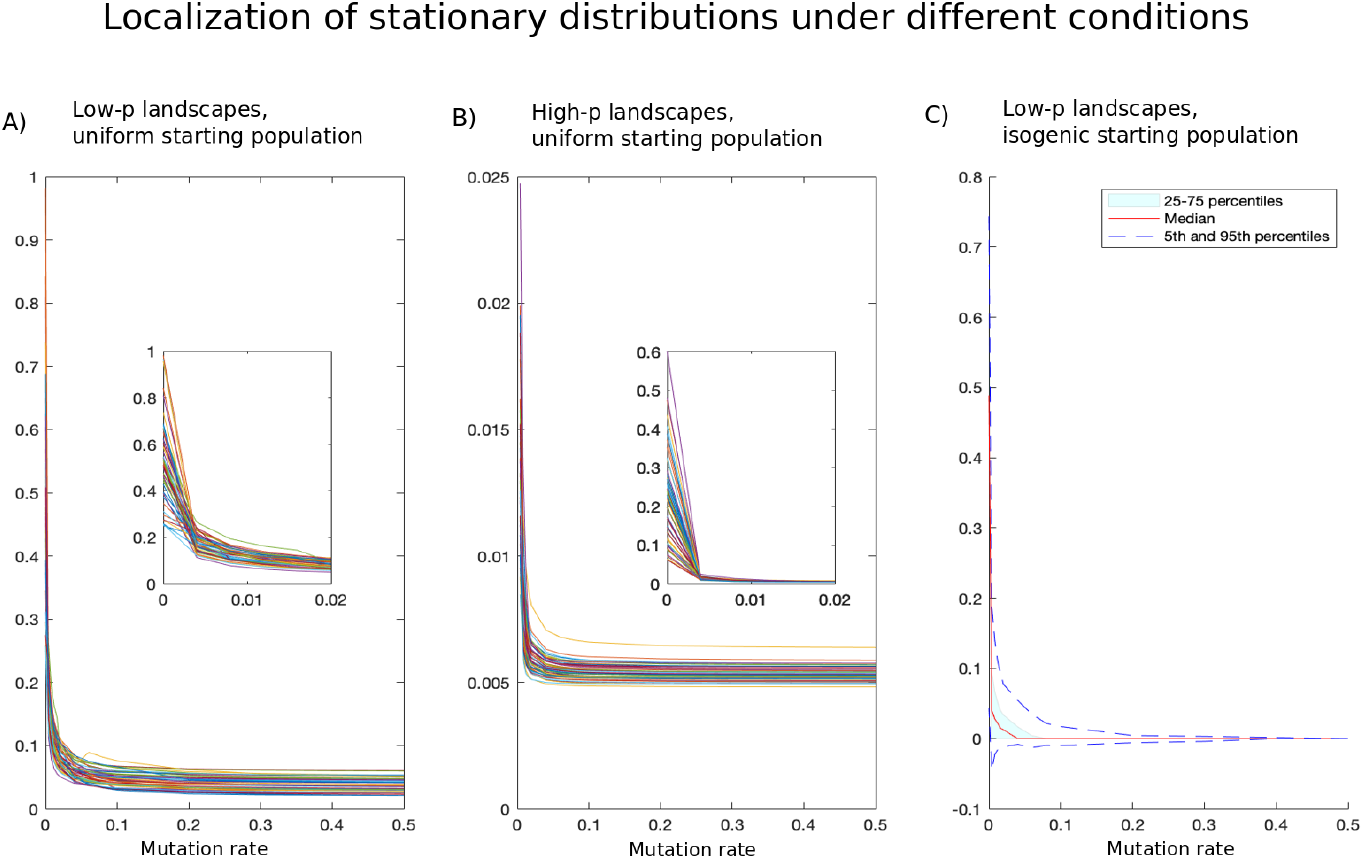
Illustrating the dependence of localization on parameters in the haploid regime with mutation and recombination. Here the rate of recombination (*r* = 0.1) is held fixed and the rate of mutation is allowed to vary. As the figure shows, when mutation is low, localization drastically increases and approaches 1. When mutation increases past a certain threshold, the localization measure stabilizes. Trajectories are plotted for different random landscapes. Similar patterns are observed for low and high *p*, but the baseline localization is higher for low *p*.

However, in low *p* landscapes (Figure 4 A) the stationary distribution’s localization is higher (as we have seen is the case for eigencentrality), and the transition to high localization occurs at a higher mutation inflection point, as compared to high *p* landscapes (Figure 4 B). Moreover, in low *p* landscapes localization tends to be, but is not always, higher when the initial population is isogenic as opposed to uniform (Figure 4 C). In high *p* landscapes, we find that the initial population distribution has a limited effect on the stationary distribution.

We next observe that, when mutation rate is high relative to recombination, the stationary distribution tends to be very close to the eigencentrality distribution, for both *p* regimes but especially for high *p* landscapes, thus affirming the importance of eigencentrality even in the presence of recombination. As mutation rate decreases, the stationary distribution’s distance to the eigencentrality distribution increases due to its increased localization.

In addition to the gradual process of increased localization, the stationary distribution may experience abrupt shifts for certain values of *µ*. These shifts can be measured by tracking the eigencentrality of the mode of the distribution (Figure S2), which at high values of *µ* is precisely the peak of eigencentrality for the vast majority of landscapes, but which abruptly falls for several landscapes as *µ* decreases, indicating sudden shifts to less eigencentral regions of the graph. These abrupt falls in the eigencentrality of the mode are more prevalent, more substantial, and begin at higher values of *µ* for low-*p* landscapes when compared to high-*p* landscapes. For the latter, even when the eigencentrality of the mode falls it is still much higher than the baseline (average) eigencentrality of nodes in the landscape.

Accordingly, the average eigencentrality of the stationary distribution reliably increases for high-*p* (Figure 5 A) landscapes as mutation decreases, due to the effect of localization – localizing around a high-eigencentrality region increases the stationary distribution’s average eigencentrality even if its mode is not at peak eigencentrality. In contrast, for the low-*p* landscapes (Figure 5 B), while we observe a general trend of increasing average eigencentrality of the stationary distribution as mutation decreases, several landscapes see a fall in stationary distribution average eigencentrality, which can be explained by increased localization around low-eigencentrality regions. For another view on the interplay of eigencentrality and localization, Figure S1 in Supplemental Information tracks the total distance between the stationary distribution and eigencentrality, a distance which can increase either due to increased localization or due to a shift in location.

**Figure 5:**
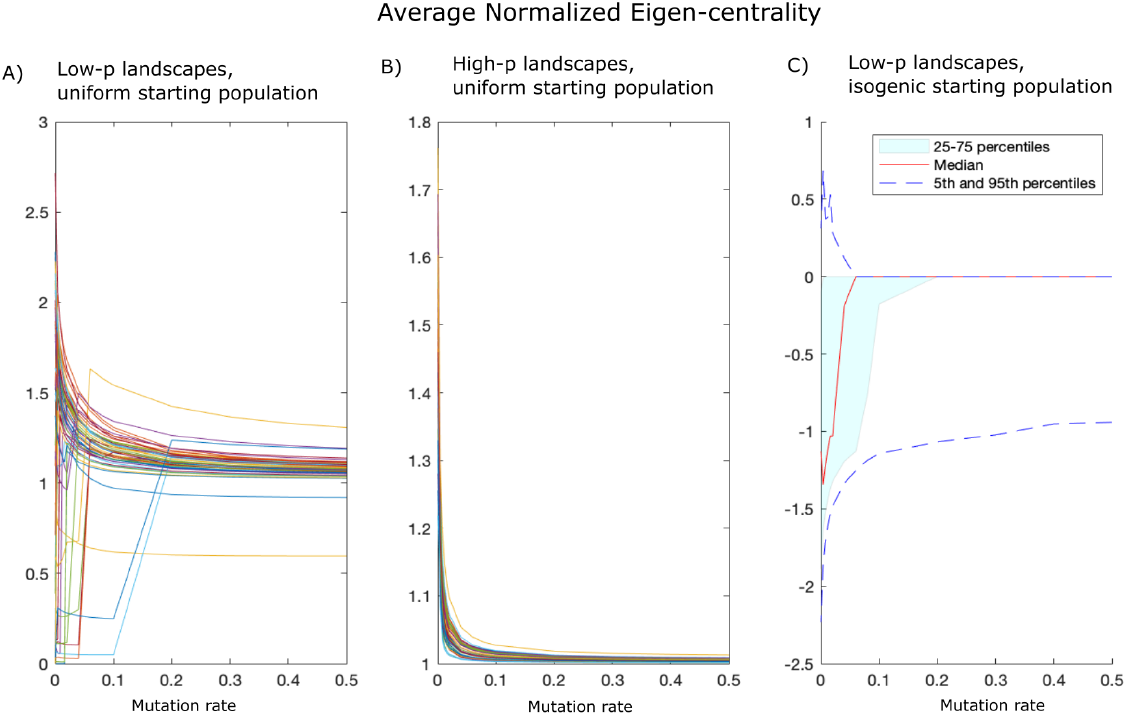
Illustrating the potential divergence between stationary distribution and eigencentrality in the presence of recombination. In A) and B), the average eigencentrality is measured for stationary distributions of 50 randomly generated landscapes (in A, with low *p* and in B with high *p*) under a uniform starting population, then normalized by dividing by the corresponding measure of the eigencentrality distribution itself. A value greater than 1 implies a population distribution which is both located along eigencentral nodes and *localized*. While in the high *p* case, average eigencentrality reliably increases with decreased mutation rate due to increased localization, in the low *p* case as we decrease mutation rate, the average eigencentrality of the stationary distribution first increases due to localization, but then for certain landscapes it sharply decreases, as recombination dominates and results in shifting the region of genotype space occupied by the population to a less eigencentral region. In C), we plot, for a given landscape the the distribution of average eigencentralities for each possible isogenic starting population, normalized so that 0 represents the average eigencentrality under uniform starting conditions. We see that the result depends on initial condition and, more often than not, results in a lower eigencentrality due to getting stuck in a low-eigencentrality region.

Under low-*p* landscapes with isogenic starting populations (Figure 5 C), most but not all stationary distributions are equally or less eigencentral in comparison to a uniform starting population, whether eigencentrality is measured by distance to the eigencentrality distribution, average eigencentrality or the eigencentrality of the mode. We interpret this to be an effect of increased local influence of the starting state of the population. Indeed, we observe that in almost all cases in which the mode of the stationary distribution differs when we change from a uniform to an isogenic starting distribution, it moves closer to the starting point of that isogenic distribution.

In the case of isogenic starting distributions, we observe a bifurcation process. For high mutation rates, under the majority of starting points, the stationary distribution for that isogenic starting population is the same as with a uniform starting population. As mutation rate lowers, however, the possible stationary distributions bifurcate into an increasing number of different possibilities, representing basins of attraction. Thus, the local influence of the starting point on the final distribution increases as mutation rate decreases relative to recombination. This reflects a transition away from eigencentrality (the only equilibrium associated to mutation dynamics) and toward the equilibria of the recombination dynamics, which can depend on initial condition. We now investigate these equilibria.

#### Stable Equilibria for Recombination

In order to appreciate the role of recombination in the dynamics of neutral evolution, it is instructive to examine the extreme case of recombination only, with no mutation. In this case, we make some observations about stable equilibria.

In our observations, in all the instances we have looked at stable equilibria of the recombination dynamics are always *closed under recombination*: that is, any two genotypes with non-zero frequency in these equilibria will produce a viable genotype when exposed to any recombination event. It remains an open question to mathematically prove this observation, or to give general conditions under which it holds. In fact, a family of genotypes closed under recombination is necessarily a sub-hypercube of the *n*-dimensional hypercube, where some alleles are fixed and the other loci may assume any combination of alleles. Thus, the process of recombination only becomes localized to a small family of this type. This gives a theoretical explanation of the observation that recombination entails higher localization, since in the extreme case of no mutation, we see localization to a closed family of viable genotypes, which is necessarily small in a typical random network.

Furthermore, we note that the equilibrium distribution on such a closed family will be of Hardy Weinberg type.

We emphasize that for each landscape there can be more than one stable equilibrium. A generic initial distribution will converge, when put through evolutionary dynamics, to one of these equilibria, depending on the inital condition. When a small mutation rate is added, in the case where recombination dominates over mutation, the population will tend to converge to a distribution which is close to one of the above-mentioned equilibria, but somewhat more diffuse due to the presence of mutation. Again, the specific attractor it converges to depends on the initial condition. In Figure 6, we plot a sample landscape (Russian roulette with *n* = 10, *p* = 0.2) and four of its basins of attraction. For each genotype, evolve to stationarity a population beginning with an initial isogenic population of that genotype. We observe that the genotypes are partitioned into 7 classes depending on which stationary distributions these isogenic initial distributions converge to. We consider these 7 stationary distributions *attractors*, and the set of genotypes which converge to a given attractor its *basin of attraction*. In Figure 6, we plot the 4 largest basins of attraction for the given landscape. Notice that each basin of attraction tends to be a connected region of the graph, which overlaps with the high-frequency nodes of its attractor, but also includes several low-frequency nodes.

**Figure 6:**
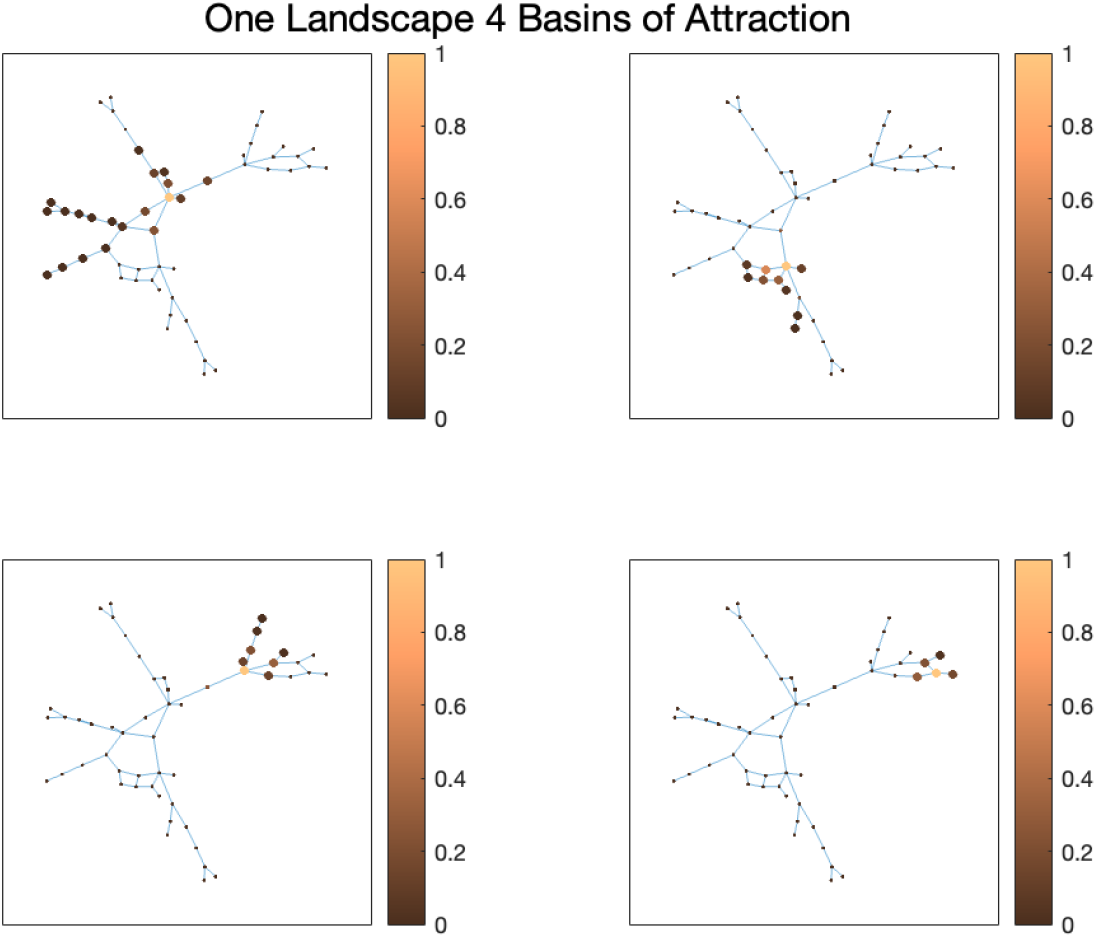
The four largest basins of attraction of the given landscape. For each panel, a node is depicted larger if it belongs to the given basin of attraction, and the color represents each node’s frequency in the attractor’s stationary distirbution.

### The Case of Diploid Population

Next, we consider a diploid population. We assume *n* loci with 2 alleles each.

We consider the following two paradigms for generating families of holey landscapes. In the first paradigm, we think of a “locus” as actually signifying a single nucleotide. Thus the “genotype” under consideration is actually a gene or even a portion of a gene. In this view, a fitness value of 1 indicates maintenance of the wild-type functionality of this gene, whereas a fitness value of 0 indicates a deleterious effect on this functionality.

In the second paradigm, we think of a “locus” as representing an entire gene. In this view, a fitness value of 1 indicates viability of the genotype as a whole.

In this paper, we to favor the first paradigm in applying our framework, since it more closely matches the model: each nucleotide position has a small number of possible values, whereas a gene has a vast number of possibilities. In order to work with our model under the second paradigm, one would have to restrict attention to only a small select set of mutants, which may be unrealistic.

As such, working in the first paradigm, we require the following symmetry: the fitness of a pair of gametes is the same regardless of the order of gametes. (In contrast, if one were working in the second paradigm, they may require a stronger symmetry: the fitness of a diploid remains the same when transposing the alleles at a specific locus).

Next, we consider different viability functions that determine neutral diploid landscapes. A first approach is analogous to the Russian roulette in the haploid case: each (unordered) pair of gametes is randomly and independently deemed viable with probability *p*. We note that the mutational network structure of *n*-locus diploids is equivalent to that of 2*n*-locus haploids. However, even in the absence of recombination, the diploid case is drastically different due to the random assortment of gametes. Indeed, the random assortment of gametes that occurs regardless of recombination can be seen as a de-facto recombination-like event. To put it concretely, diploid reproduction with mutation but no recombination is very similar to – though not quite the same as – haploid reproduction with mutation and a single recombination event at a 100% rate (the reason that these two are different is that diploid recombination is indifferent to which gamete is picked).

Due to our understanding of haploid recombination, which dominates over mutation when the recombination rate is high and drives the population away from eigencentrality, we would expect the same to hold for diploid populations even in the absence of explicit recombination. Indeed, we observe that, regardless of recombination and even with a high mutation rate, the population does not approach high eigencentrality. Indeed, in the Figure 7 A we plot average eigencentrality of the stationary distributions of 50 random landscapes under the diploid dynamics with no recombination, only random assortment (normalized so that 1 represents the average eigencentrality of the eigencentrality distribution itself). The figure shows that the average eigencentrality widely varies among landscapes, with no clear trend (compare to the haploid case). On the other hand, increased mutation does still lead to consistently lower localization by spreading out the population (figure 7 B).

**Figure 7:**
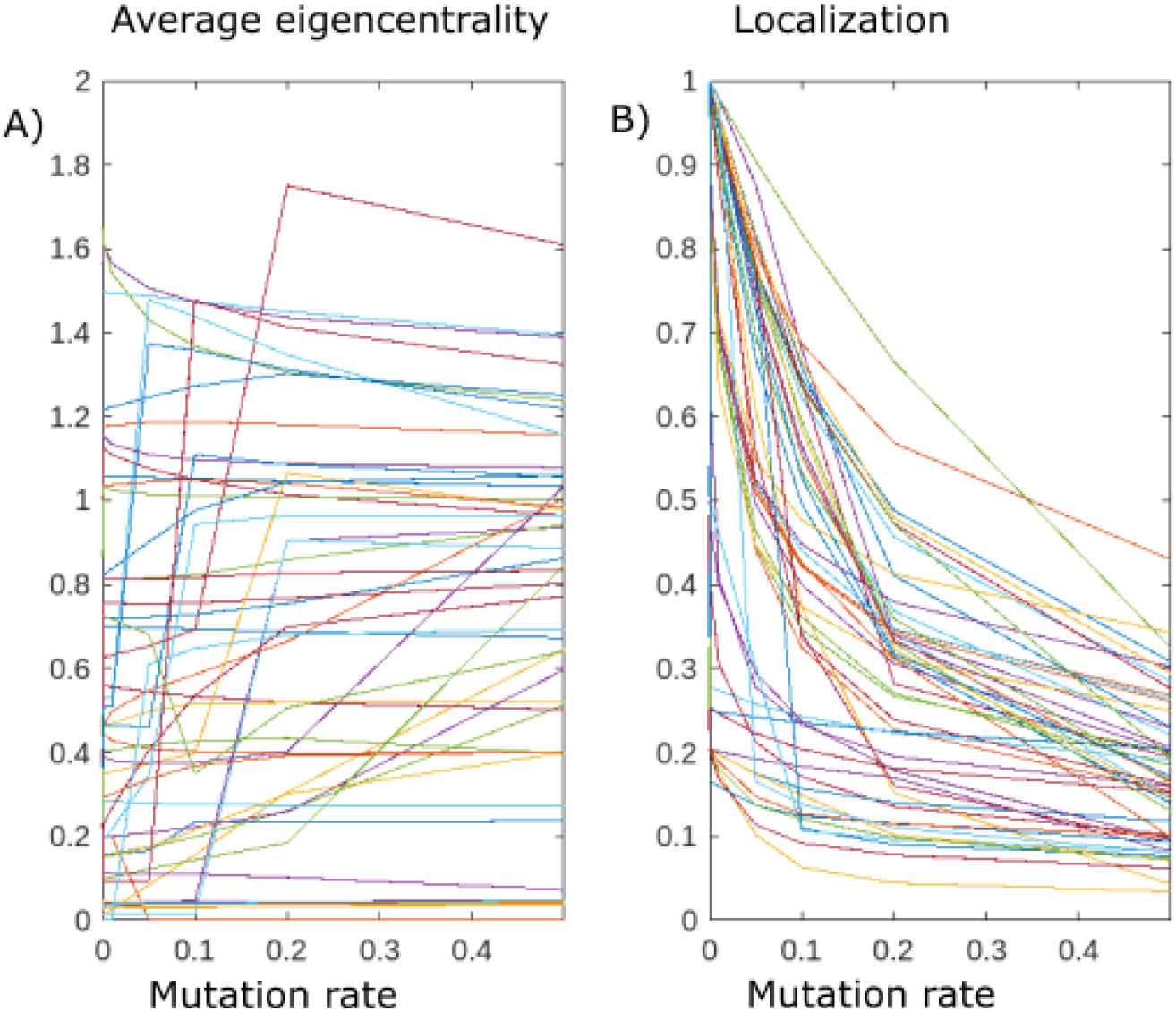
A) Average Eigencentrality (normalized so that 1 is the average eigencentrality of the eigencentrality distribution itself) and B) Localization for 50 random diploid (unstructured) Russian roulette landscapes.

Adding in recombination (crossing over) does not have a clear effect on the distribution or its localization. The stationary distributions are largely similar with or without recombination. In the high-mutation case, 90% of the landscapes had the same mode when recombination was added, and in the zero-mutation case, 80% of the landscapes had the same mode.

An alternative approach is to define the viability of a genotype as a function of the individual gametes’ viabilities. Here, we consider two possible ways of combining a viability function on gametes to form a viability function on diploids: dominance of lethality, requiring that both gametes be viable in order for the genotype to be viable; and recessiveness of lethality, requiring that at least one gamete be viable in order for the genotype to be viable.

In the case of dominance, we note that under the dynamics of mutation and recombination, the *gamete* distribution evolves independently in a manner nearly identical to the haploid dynamics, and the diploid distribution is simply the product of the gamete distribution. Indeed, when we defined our model of diploid recombination dynamics, we noted that the recombination tensor for diploid distributions behaves as the product of corresponding haploid recombination tensors for the individual gamete distributions. The same is not quite true for the mutation dynamics, due to our simplifying modeling assumption that there can only be at most one mutation in every generation. Thus, a diploid under our model can be exposed to at most one mutation per generation whereas two independent haploids could be exposed to two. However, as mutation rate is generally small, the chance of two mutations can be treated as negligible. Therefore, the diploid mutation dynamics is approximately the same as independently exposing the gametes to haploid dynamics and then taking the product distribution (we could change our model to make this approximation exact, but currently choose not to for simplicity). Finally, the selection dynamics for diploids also behave according to independent selection of gametes, by virtue of the underlying assumption of dominance of lethality which guarantees a diploid is viable exactly when both its gametes are. Therefore, the entire evolutionary dynamics behave approximately as the product of haploid dynamics of the gametes. Indeed, simulations show that, for a diploid landscape with dominance of lethality of size 100, the difference between the stationary distribution of the diploid dynamics and the product of haploid dynamics is very small: a difference of less than 10^−4^ for each diploid genotype.

Due to the above theoretical observation, we do not analyze and simulate this case further as it is approximately reducible to the haploid dynamics already mentioned. In particular, in the mutation-only case we see that the population distribution will converge to the product of the eigencentrality distributions of the individual gametes. This product is in fact identical to the eigencentrality distribution of the diploid landscape itself (see Supplemental Information on product structures for more detail).

In the case of recessiveness of lethality, the diploid stationary distribution, even in the mutation-only case, does not have such a simple description, and is not given in terms of the eigencentrality of the gametes. Indeed, in a recessive model, a viable diploid can have a gamete which is on its own inviable, and thus does not have a notion of eigencentrality.

For these landscapes, unlike the Russian Roulette diploid landscapes, we observe that the stationary distribution *does* tend to closely align with its eigencentrality. We first consider the case of only mutation, and no recombination (only the random assortment of gametes). Figure 8 shows relevant statistics for a typical landscape:

**Figure 8:**
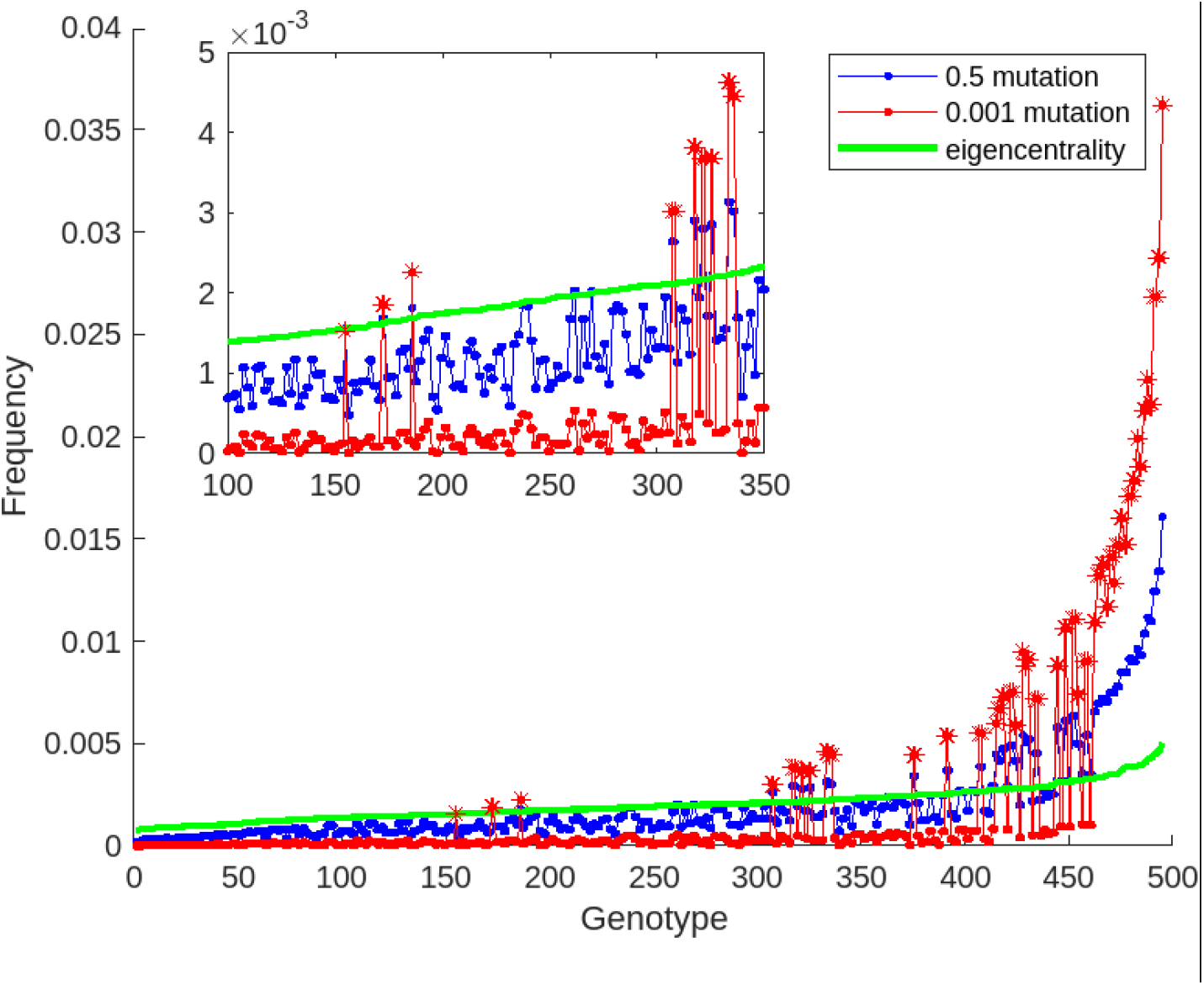
For a fixed landscape in the recessive lethality family of diploid landscapes, we calculate stationary distributions under high mutation (0.5) and low mutation (0.001). In both cases there is no explicit recombination, only random assortment of gametes. The blue and red lines show the stationary distributions of the evolutionary dynamics under the two cases, respectively. Genotypes are sorted in ascending order of their eigencentrality measure, and eigencentrality is plotted in the solid green line. The insert at the top left zooms in on a portion of the plot. Asterisks are placed over those genotypes both of whose gametes are viable.

In Figure 8, we order the landscape’s genotypes in order of increasing eigencentrality (whose value is indicated by the green line), while the blue and red lines indicate the stationary distributions under high (0.5) and low (0.001) values of mutation, respectively. The asterisks are placed along those genotypes where both gametes are viable. We can make several observations: first, we notice that the eigencentrality distribution is characterized by low localization– the distribution is spread out over all genotype space. Moreover, eigencentrality itself tends to reward those double-viable genotypes, which makes sense given that those genotypes will tend to have many more viable neighbors as any single mutation preserves the viability of the diploid. However, notably there are a few exceptions to this pattern: some double-viable genotypes have lower eigencentrality than some single-viables. This can be explained by the fact that eigencentrality rewards not just the number of viable neighbors, but those neighbors’ centrality as well. Thus, a single-viable in a central region of the graph can be rewarded over a double-viable whose location in the graph is less central. Next, we can observe that, especially in the higher-frequency genotypes of the distributions, there is a strong relationship between their ranking in eigencentrality and their ranking in frequency, confirming that the stationary distribution aligns with eigencentrality. We observe that the key effect of decreasing mutation rate is to boost the frequency of the double-viables relative to the single-viables. However, within each of these pools the relative relationship of frequencies remains similar, as can be seen in the insert, comparing the red line and blue line among the single-viable genotypes. This effect of boosting the double-viables can be understood by considering the limit case when mutation rate is brought down to 0. In this case, it can be shown mathematically (see Supplemental Info) that only the diploids where both gametes are viable survive in the stationary distribution, at a proportional frequency to their initial distribution.

Relatedly, decreasing mutation leads to increased localization of the stationary distribution, which is consistent with the tendency of the distribution toward the small set of double-viable genotypes. If we think of random assortment as a type of de-facto recombination, then this behavior is also consistent with the dynamics we’ve observed of landscapes under recombination, tending toward higher localization when mutation is low. Specifically, the set of double-viables acts as an attractor which is closed under the random-assortment operator.

Next, we consider the case in which recombination is added in addition to mutation. Here we observe the same broad patterns as above: the stationary distribution tends to align with eigencentrality (although there are now more instances of breaking away from eigencentrality when mutation is decreased), and localization increases as the mutation rate decreases. Moreover, the stationary distribution is more localized when compared to the analogous stationary distribution with no recombination but the same mutation rate.

Given the importance of eigencentrality to the stationary distributtions of these recessive lethality landscapes, it is an interesting question to relate the eigencentrality of a diploid genotype to its component gametes and their position within the fitness landscape of gametes. One preliminary observation is that, while for the most part the eigencentrality favors diploids where both gametes are viable, nonetheless some inviable gametes have a higher total frequency in the eigencentrality distribution than some viable gametes. This can be explained by the observation that some inviable gametes may still have a high number of highly eigencentral viable gametes, which will make them more favored when compared with viable gametes that do not have many viable or central neighbors. This observation reinforces the importance of position within the network, which can in some cases prove more important than a gamete’s immediate fitness. However, as explained above, the frequency of such inviable gametes is further lessened as the mutation rate is decreased.

## Discussion

To summarize, our study details and examines the richness of phenomena associated with neutral evolution of large populations, with an emphasis on the importance of *network structure* in shaping these evolutionary dynamics, while also examining different regimes in parameter space that lead to qualitatively distinct evolutionary behaviors. Our findings highlight the complex interplay between mutation, recombination, and the structure of fitness landscapes in shaping evolutionary trajectories, even in the absence of direct selection. They underscore the importance of considering the global structure of these landscapes to understand evolutionary outcomes.

Two properties of population distributions that we pay particular attention to are *eigencentrality*, referring to the location of the distribution among genotypic nodes which can be viewed as more central (or robust) in the network, and *localization*, a measure of the homogeneity of a distribution, where most of the population is concentrated along a small region of genotypic space.

Though knowledge of these concepts’ role in neutral evolution is not new, we believe that they remain under-explored, and a full understanding of the range of implications is still lacking. Towards a more robust understanding, we carry out a systematic study of these properties under different regimes in parameter space and their dependence on network structure.

To summarize our findings broadly: under a regime of only mutation in haploid populations, the population distribution converges to the eigencentrality distribution regardless of mutation rate or initial conditions. This eigencentrality distribution may or may not be localized depending on the network structure: for random landscapes the localization increases as the probability of viability decreases. Once recombination is added to the evolutionary dynamics, these dynamics become more complex: they depend both on the initial conditions and on the balance between the parameters. Broadly speaking, there are two regimes, with a phase shift between them depending on the balance between mutation and recombination rates: if mutation dominates, then the stationary distribution resembles eigencentrality, while being more localized due to the influence of recombination; whereas if recombination dominates, then the distribution no longer resembles eigencentrality but instead approaches attractors of the recombination dynamics, where the specific attractor that the distribution goes toward depends on initial conditions. These attractors tend to be very localized, explaining the tendency of recombination to increase the distribution’s localization. We note that while [21] made the general observation that the presence of recombination tends to increase average population robustness, we further complicate this observation, as high recombination can often lead a population away from the most eigencentral region of the graph, while on the other hand compensating for this by increasing the population’s localization. Finally, diploid populations present a different case. Random diploid landscapes, even without explicit recombination, behave qualitatively similarly to haploid landscapes where recombination dominates – that is, they bear no relation to eigencentrality. However, if the diploid landscape is *structured*, such that diploid viability is a function of the component gametes’ viabilities, then in fact the stationary distribution does bear a close relationship to eigencentrality, showing that eigencentrality has the potential to be a relevant factor in the neutral evolution of diploid populations as well.

The potential broader implications of our findings are myriad. We suggest that the role of neutral evolution should be taken seriously as a possible explanation of phenomena that are typically attributed to selection. We note that, while the neutral landscapes we study do not have *direct* selection in the sense that there is no fitness differential between viable genotypes, there is still a kind of indirect selection involving the location of a genotype within the network, and this type of indirect selection can give rise to similar phenomena that one sees with selection, albeit in a more subtle form.

For example, conserved regions within the genome may not necessarily reflect a selective advantage. Instead, they may be the result of neutral evolutionary dynamics in which recombination rates predominate, which can yield very high population localization for the neutral landscape of that portion of the genome, even without direct selection. Similarly, in cases where separate populations or even distinct species appear to exhibit evolutionary convergence, it may not be accurate to attribute this convergence solely to selective pressures favoring a particular sequence. Such convergent evolution might instead be a manifestation of the inherent structure of the neutral fitness landscape. This could be consistent with neutral evolution in either the mutation-dominant or recombination-dominant regimes, where neutral evolutionary processes guide different initial populations toward similar genotypic regions without the direct influence of selection.

When analyzing the patterns of evolutionary trajectories, it is essential to recognize that different regions of the genome may not evolve in qualitatively identical ways. Our study suggests that the qualitative properties of these evolutionary trajectories are highly dependent on the structure of a particular genetic site’s fitness landscape, or even on the relative rates of mutation and recombination which can vary over the genome. This highlights the need for a nuanced approach to studying evolutionary processes, where the specific dynamics of mutation and recombination as well as structural characteristics of the fitness landscape are carefully considered before drawing conclusions about the presence and relevance of selection in shaping these trajectories [24, 25, 26, 27, 6].

The fact that qualitatively distinct evolutionary dynamics can arise depending on the balance between mutation and recombination also has potential implications, which we have not explored here, on the evolution of recombination and the regulation of these rates.

Many questions remain which can be tackled in future work. We assumed for simplicity an infinite population and a binary (0 or 1) fitness function. It is understood that relaxing the fitness function so that deleterious mutations do not lead to 0 fitness but rather to a sharp drop in fitness, does not significantly alter the general evolutionary dynamics [16], but a more thorough testing of this understanding is merited. Furthermore, the study of finite (even if large) populations adds the potential of random drift as an additional complicating factor, whose interaction with the dynamics we’ve identified requires further study.

Moreover, there is potential in applying mathematical tools from random graph theory and spectral graph theory to lend a theoretical foundation to the observations we’ve made.

Finally, our simulations could be augmented by considering multiple populations with a migration rate or adding a spatial component, which can shed light on the implications of neutral evolution on speciation and the maintenance of genetic divergence between populations despite gene flow. Typically, the focus has been on selective pressures—such as disruptive selection or clines—that drive populations toward distinct evolutionary paths, or else on the emergence of genetic incompatibilities. However, future work could examine these phenomena through the framework of neutral evolution. Specifically, as we have seen, recombination dynamics can drive populations to different attractors depending on the initial conditions. We hypothesize that these attractors, even if not fully incompatible with each other, could in some cases exert enough pull that distinct populations at different attractors can maintain their divergence even in the presence of limited amounts of gene flow between them.

## Supplemental Info

### Further details on Proposition 1

The limit of **p**_*t*_ as *t* → ∞ can be analyzed using the eigenvectors of *B*. Since *B* is symmetric, it is diagonalizable. Thus any vector **p**_0_ can be decomposed **p**_0_ = Σ_*i*_ *v*_*i*_ as a sum of eigenvectors *v*_i_ with eigenvalues *λ*_i_, so that 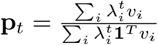.

As *t* grows large, the terms associated with highest-magnitude eigenvalues dominate. In fact, as *t* approaches infinity, **p**_*t*_ will always converge to the same stationary distribution, regardless of the initial distribution **p**_0_. This is a consequence of the Perron-Frobenius Theorem applied to *B*.

Indeed, since *B* has non-negative entries and is irreducible (meaning its underlying graph is connected, which follows from our assumption that *G’* is connected), Perron-Frobenius guarantees that the spectral radius *ρ*(*B*) is an eigenvalue of *B* whose eigenspace has dimension 1; moreover, we can choose an eigenvector with eigenvalue *ρ*(*B*) all of whose entries are positive, and conversely any eigenvector all of whose entries are positive must have eigenvalue *ρ*(*B*). If we normalize by the requirement that the entries add to 1, there is a unique such vector, which we will denote 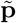. We also note that −*ρ*(*B*) is not an eigenvalue of *B*. Indeed, this is again guaranteed by Perron-Frobenius, as a consequence of the fact that *B* has period 1 (meaning the greatest common divisor of the lengths of loops in the graph underlying *B* is 1; this follows automaticaly from the fact that *B* has positive entries on the diagonal).

Now, we claim that **p**_*t*_ approaches 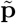 as *t* → ∞, no matter the initial distribution **p**_0_. The key is that the eigenvector decomposition of **p**_0_ is bound to include a nonzero component along 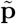. Indeed, if this were not the case, let *r < ρ*(*B*) be the highest magnitude of the eigenvalues in the eigen-decomposition of **p**_0_.

Since eigenvalues with magnitude *r* (that is, *λ* = *±r*) dominate the rest, **p**_*t*_ will approach an oscillation of the form *v ± w*, where *v* and *w* are eigenvectors with eigenvalues *r* and −*r*, respectively. So the average of **p**_*t*_ and **p**_*t*+1_ approaches *v*, an eigenvector with eigenvalue *r*. But the **p**_*t*_ have all positive entries, so by Perron-Frobenius, *v* must equal 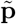, contradicting *r < ρ*(*B*).

In fact, the stationary distribution 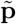 has a simple interpretation. Note that, due to the relationship 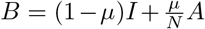, the matrices *A* and *B* have the same eigenvectors. Moreover, though they do not have the same eigenvalues, there is an order-preserving one-to-one correspondence between the eigenvalues of *A* and *B*. Thus, the eigenvector 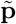 is also the eigenvector of *A* with largest eigenvalue. Since *A* is the adjacency matrix of the graph *G ’*, the vector 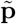 is also known as the eigenvector-centrality of *G ’*. That is, in the stationary distribution, the nodes *v* ∈ *V ’* are distributed according to their eigencentrality in the graph *G ’*. Moreover, we note that, in our model, the stationary distribution does not depend on the mutation rate *µ*. The rate *µ*, however, does affect the rate of convergence to the stationary distribution.

### Proof of eigencentrality on interval

We claim that the eigencentrality of a geometric interval {1, …, *n*} is 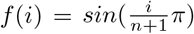 for 1 ≤ *i* ≤ *n*. Thanks to Perron-Frobenius, since this function is strictly positive, we can prove that it is indeed the stationary distribution by proving that it is an eigenvector of the adjancency matrix *A* given by *A*_*ij*_ = 1 if and only if |*i* − *j*| = 1.

This follows from simple trigonometry. We claim *f*, restricted to 1 ≤ *i* ≤ *n*, is an eigenvector of the matrix *A* with eigenvalue 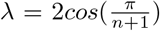. Indeed, the eigenvector equations boil down to

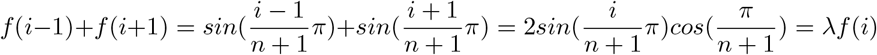

Note that this captures the boundary conditions as well, since *f* (*i*) = 0 when *i* = 0 or *n* + 1.

The above equations are a consequence of the general trigonometric identity

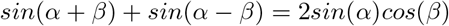

for any angles *α, β*.

The formula for the eigencentrality of a geometric box can then be obtained by reducing to the case of an interval, by considering product structures of landscapes, discussed below.

### Product Structures

We now focus on the special case in which a fitness landscape can be decomposed as a product of fitness landscapes. We formalize this notion below.

Consider two fitness landscapes: the first, a graph *G*_1_ = (*V*_1_, *E*_2_) together with a fitness function *f*_1_ : *V*_1_ → ℝ^≥0^; and the second, a graph *G*_2_ = (*V*_2_, *E*_2_) together with a fitness function *f*_2_ : *V*_2_ → ℝ^≥0^.

We define the product of these two fitness landscapes to be the following. Its vertex set is *V*_1_ *× V*_2_, and its edges are given by 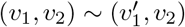 whenever *v*_1_ is a neighbor of 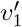 in *E*_1_, and 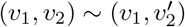 whenever *v*_2_ is a neighbor of 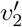 in *E*_2_. In other words, the neighbors of (*v*_1_, *v*_2_) are the results of substituting one of *v*_1_, *v*_2_ with a neighbor, and leaving the other one fixed. Finally, the fitness function on the product is given by *f* (*v*_1_, *v*_2_) = *f*_1_(*v*_1_)*f*_2_(*v*_2_).

In the special case of holey landscapes, where *f*_1_ and *f*_2_ only take on values of 0 or 1, then this means *f* (*v*_1_, *v*_2_) = 1 whenever both *f*_1_(*v*_1_) = 1 and *f* (*v*_2_) = 1, and *f* (*v*_1_, *v*_2_) = 0 otherwise.

The structure described above can be generalize to *n*-dimensions, representing a genetic system composed of *n* genes whose alleles can mutate among *m*_*i*_ neighboring alleles the *i*’th gene can assume. Such a product structure as arising from a genetic system that have no epistatic interaction, in the sense that a fatal mutation in the one gene leads to inviability regardless of the genetic background of the other set of genes.

In the case of holey landscapes, we now demonstrate that the stationary distribution of the product *V*_1_ *× V*_2_ under the setup above is simply the product of the individual stationary distributions on *V*_1_ and *V*_2_. Let 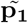 and 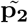 denote the stationary distributions on *V*_1_ and *V*_2_, respectively, and let 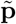 denote the stationary distribution on the product.

To show that 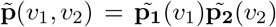, it suffices by Perron-Frobenius to show that 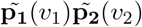 forms an eigenvector for the adjacency matrix of viable nodes in *V*_1_ *× V*_2_. Indeed, it has eigenvalue *λ*_1_ + *λ*_2_, where *λ*_1_ is the eiegenvalue associated to 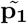 in *V*_1_ and *λ*_2_ is the eigenvalue associated to 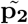 in *V*_2_. This boils down to the following set of equations, for each pair of viable nodes 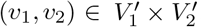:

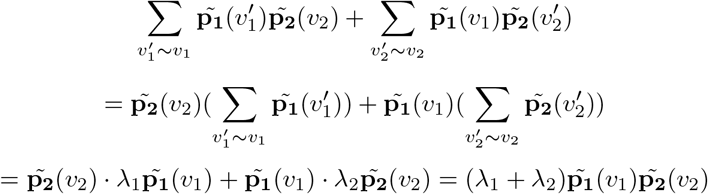

Thus, given such a product structure *V* = *V*_1_ *× V*_2_, we may reduce the study of the stationary distribution to the independent components *V*_1_, *V*_2_.

### Localization of eigencentrality on an interval

Given our formula above for the eigencentrality on an interval {1, …, *n* − 1}, if we normalize eigencentrality so it sums to 1, the formula becomes:

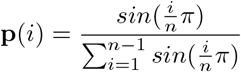

The normalized Simpson diversity index of the eigencentrality on an interval {1, …, *n* − 1} is given by *n***p** · **p** =

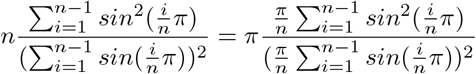

Both the numerator and denominator above can be understood as Riemann sums, so that in the limit the expression becomes

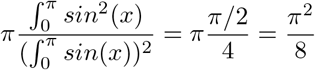

as claimed.

A similar calculation can be carried out in the case of a geometric square.

### Diploid landscapes with recessive lethality: the no mutation, no recombination case

In this section we provide a proof for the claim that, in diploid landscapes that are generated using recessive lethality, the dynamics in the case of no mutation and no recombination (that is, only random assoortment of gametes) leads to a distribution in which only the double-viable diploids are represented, with viable gamete frequencies proportional to their frequencies in the initial distribution. To see this, consider a diploid distribution 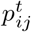, where *i* and *j* index the two gametes and the superscript *t* tracks the generation. We can assume that this distribution is symmetric, i.e. 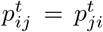 since this will be the case after one generation. Let 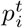, denote the total frequency of gamete *i*, so 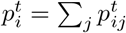. Upon recombination, we have the following identities:

If *i* is viable, then the new frequency of gamete *i* is given by

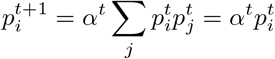

where *α*^*t*^ is some fixed re-normalization rate (to get the distribution to add up to 1). On the other hand, if *i* is inviable, then the new frequency of gamete *i* is given by

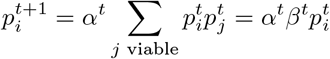

where 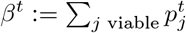. By summing 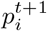 over all viable gametes, we get that

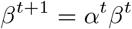

Note also that *α*^*t*^ ≥ 1 for all *t* by virtue of being a normalization factor, and *β*^*t*^ ≤ 1 for all *t* by virtue of being a probability. Thus, the sequence of values of *β*^*t*^ is increasing and bounded above, so it must converge. Thus *α*^*t*^ must converge to 1, which implies that in the limit, no inviable diploids are created upon recombination, and so in the limit all gametes are viable. Thus, the distribution of inviable gametes goes to 0, while that of viable gametes is proportional to their initial distributions, as they are always multiplied by the same factor in any given generation.

### An alternative measure of divergence from eigencentrality

An alternative perspective on the interplay between eigencentrality and localization is provided in Figure S1, showing the overall distance between the stationary distribution and eigencentrality. This distance can increase either as a result of heightened localization or due to a shift in the location of the population’s genetic makeup.

**Figure S1:**
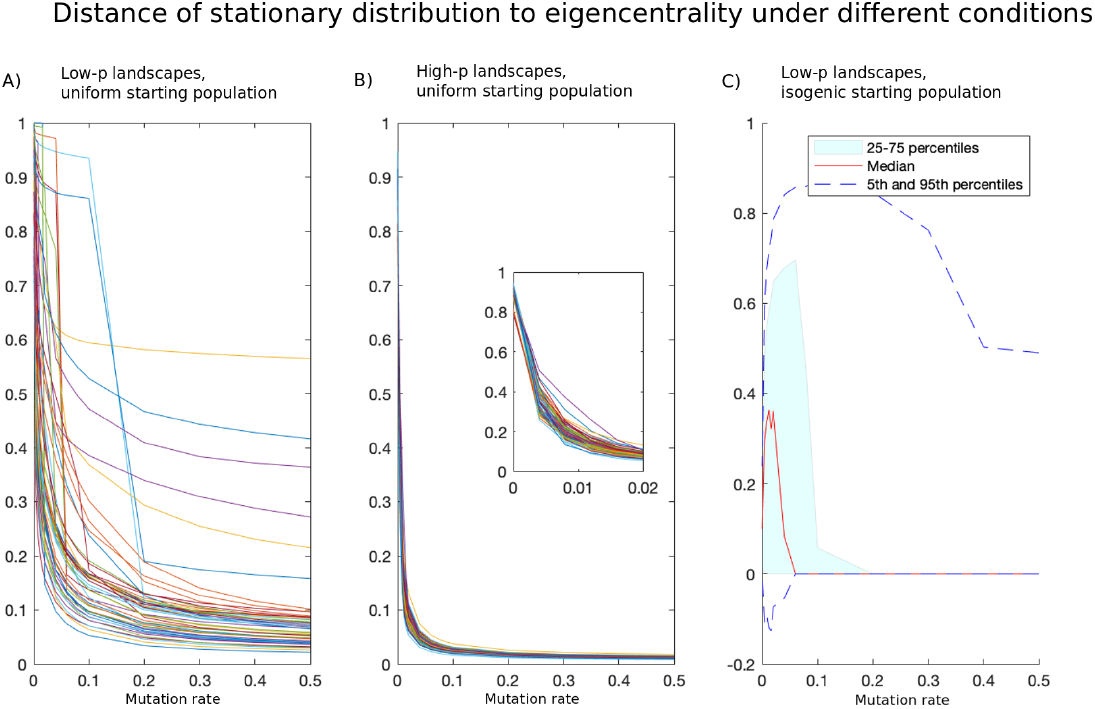
Distance to Eigencentrality

### Eigencentrality of the mode

To track the shift’s in population distribution’s locations in the network as the parameters change, it helps to consider the eigencentrality of the mode.

**Figure S2:**
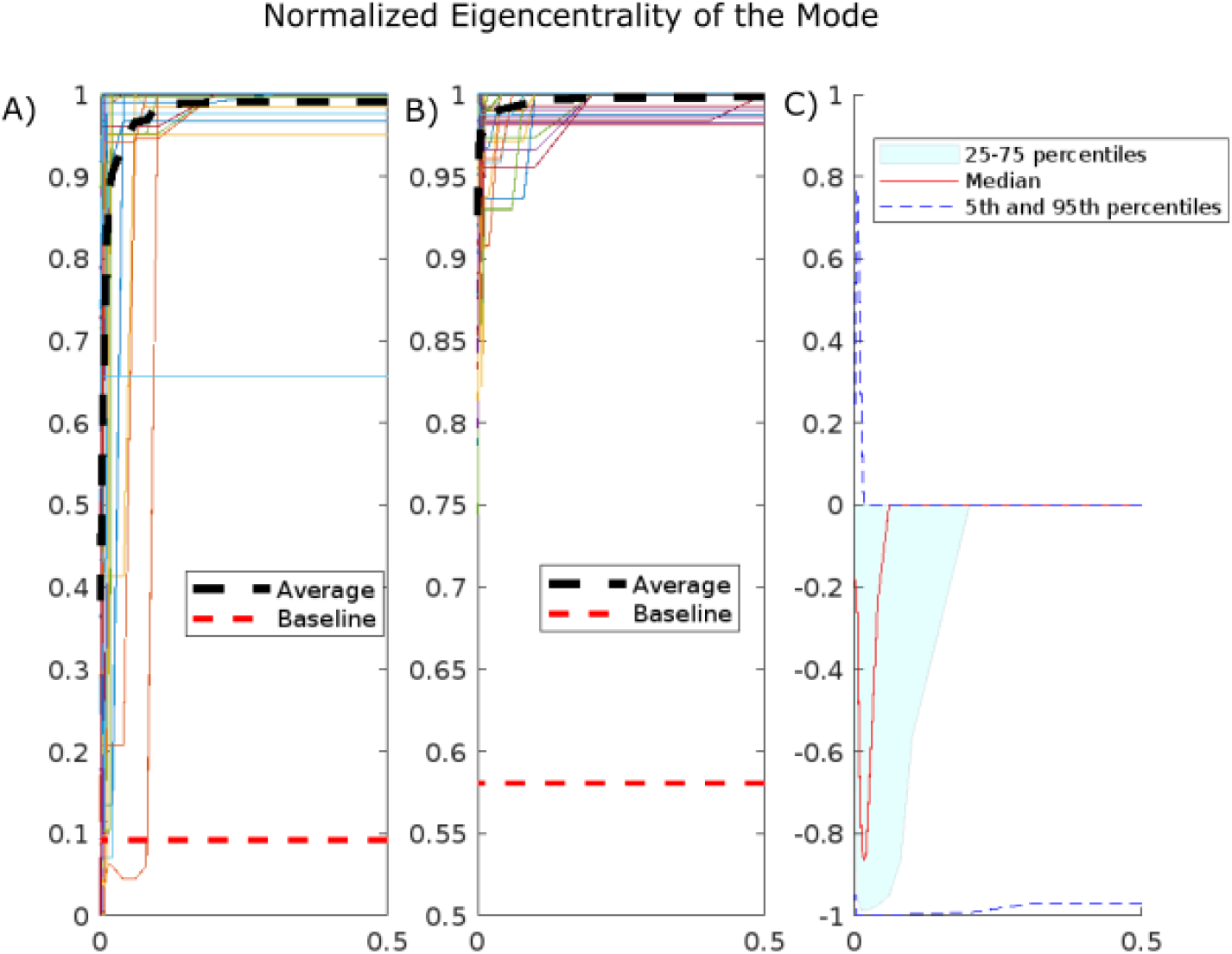
Eigencentrality of the mode, normalized so that 1 is the maximum possible value. A) low-p landscapes, with uniform starting distribution. B) high-p landscapes, with uniform starting distribution. C) low-p landscapes, with isogenic starting distribution. The black dotted lines in A) and B) track the average eigencentrality of the mode across the 50 landscapes, while the red dotted lines indicate the baseline eigencentrality (which is the average eigencentrality over the entire genotype space). The *y* axis in C represents the distribution of differences of measures: the measure for the isogenic initial population minus the measure for the uniform initial population. We see that, while for most landscapes and starting conditions the eigencentrality of the mode remains the same whether the initial distribution is isogenic or uniform, for many initial conditions the isogenic distributions lead to lower eigencentrality of the mode.

## Notes

### Competing Interest Statement

The authors have declared no competing interest.

